# Integrated ‘omic’ analyses provide evidence that a *Ca*. Accumulibacter phosphatis strain performs denitrification under micro-aerobic conditions

**DOI:** 10.1101/386516

**Authors:** Pamela Y. Camejo, Ben O. Oyserman, Katherine D. McMahon, Daniel R. Noguera

## Abstract

The unique and complex metabolism of *Candidatus* Accumulibacter phosphatis has been used for decades for efficiently removing phosphorus during wastewater treatment in reactor configurations that expose the activated sludge to cycles of anaerobic and aerobic conditions. The ability of Accumulibacter to grow and remove phosphorus during cyclic anaerobic and anoxic conditions has also been investigated as a metabolism that could lead to simultaneous removal of nitrogen and phosphorus by a single organism. However, although phosphorus removal under cyclic anaerobic and anoxic conditions has been demonstrated, elucidating the role of Accumulibacter in this process has been challenging, since experimental research describes contradictory findings and none of the published Accumulibacter genomes show the existence of a complete pathway for denitrification. In this study, we use an integrated omics analysis to elucidate the physiology of an Accumulibacter strain enriched in a reactor operated under cyclic anaerobic and micro-aerobic conditions. The reactor’s performance suggested the ability of the enriched Accumulibacter (clade IC) to simultaneously use oxygen and nitrate as electron acceptors under micro-aerobic conditions. A draft genome of this organism was assembled from metagenomic reads (hereafter referred to as Accumulibacter UW-LDO-IC) and used as a reference to examine transcript abundance throughout one reactor cycle. The genome of UW-LDO-IC revealed the presence of a full denitrification pathway. The observed patterns of transcript abundance showed evidence of co-regulation of the denitrifying genes along with a *cbb_3_* cytochrome, which is characterized as having high affinity for oxygen, thus supporting the hypothesis that UW-LDO-IC can simultaneously respire nitrate and oxygen. Furthermore, we identified an FNR-like binding motif upstream of the coregulated genes, suggesting transcriptional level regulation of the expression of both denitrifying and respiratory pathways in Accumulibacter UW-LDO-IC. Taken together, the omics analysis provides strong evidence that Accumulibacter UW-LDO-IC simultaneously uses oxygen and nitrate as electron acceptors under micro-aerobic conditions.

**IMPORTANCE:** *Candidatus* Accumulibater phosphatis is widely found in full-scale wastewater treatment plants, where it has been identified as the key organism for biological removal of phosphorus. Since aeration can account for 50% of the energy use during wastewater treatment, micro-aerobic conditions for wastewater treatment have emerged as a cost-effective alternative to conventional biological nutrient removal processes. Our study provides strong genomics-based evidence that Accumulibacter is not only the main organism contributing to phosphorus removal under micro-aerobic conditions, but also that this organism simultaneously respires nitrate and oxygen in this environment, consequently removing nitrogen and phosphorus from the wastewater. Such activity could be harnessed in innovative designs for cost-effective and energy-efficient optimization of wastewater treatment systems.

## INTRODUCTION

*Candidatus* Accumulibacter phosphatis (hereafter referred to as Accumulibacter) is the main microorganism removing phosphorus (P) in many wastewater treatment plants performing enhanced biological phosphorus removal (EBPR) (1-4). This uncultured polyphosphate accumulating organism (PAO) fosters a unique and complex metabolism that responds to changes in the availability of carbon, phosphorus, and oxygen. Under anaerobic conditions, Accumulibacter takes up volatile fatty acids (VFA) present in the wastewater and stores the carbon from these simple molecules intracellularly as poly-β-hydroxyalkanoate (PHA), while hydrolyzing intracellular polyphosphate (polyP) to phosphate, which is then released from the cell to the liquid phase (5). The subsequent addition of oxygen into the bulk liquid triggers the use of stored PHA molecules to generate energy for growth, concomitant with the uptake of phosphate from the medium to form polyphosphate, eventually leading to the efficient removal of P from the wastewater.

Analysis of the Accumulibacter lineage has led to the discovery of multiple genome variants. Using the polyphosphate kinase (*ppk1*) gene as a phylogenetic marker, Accumulibacter variants have been subdivided into two types and 14 different clades (types IA-E and IIA-I) (6-9). This genomic divergence may be responsible for observed phenotypic variations of EBPR under different environmental conditions (10-14). Among these differences, Accumulibacter’s fitness for anoxic respiration is a topic of much debate since published studies have presented contradictory findings on whether Accumulibacter can respire nitrogenous compounds. While several studies predicted that strains belonging to Accumulibacter type I could use nitrite and/or nitrate as electron acceptors (10, 15-17), other studies concluded that type I is not capable of anoxic nitrate respiration (18, 19). These studies have used different methods for clade classification, with some of them describing Accumulibacter at the type level and others at the clade level, as defined based on *ppk1* phylogeny (6). Therefore, it remains uncertain whether individual clades exhibit a consensus phenotype regarding respiration of nitrogenous compounds. It is also possible that differences in the metabolic potential of Accumulibacter may vary among strains within the same clade. Uncovering the metabolic traits characterizing distinct Accumulibacter populations will provide a better understanding of the ecological role of each of these clades/populations and the biotechnological potential of this lineage in novel nutrient removal processes.

In a previous study, we characterized the clade-level population of Accumulibacter in a biological nutrient removal reactor operated under cyclic anaerobic and micro-aerobic conditions and evaluated the ability of the enriched population to use multiple electron acceptors (8). Experimental evidence from this study led to the hypothesis that a particular clade of Accumulibacter (clade IC) could use oxygen and nitrate as electron acceptors (8) when the system is operated with cyclic anaerobic and micro-aerobic conditions. In this study, we use a combination of omics techniques to further investigate the genomic potential, gene expression, and transcriptional regulation of an enriched clade IC Accumulibacter population to further elucidate the metabolic capabilities of this species-like group. This analysis provides strong evidence that the enriched Accumulibacter clade IC population simultaneously uses oxygen and nitrate as electron acceptors under micro-aerobic conditions, and therefore, that this organism contributes to the simultaneous removal of nitrogen and phosphorus from wastewater.

## >MATERIAL AND METHODS

### Operation of Lab-Scale Sequencing Batch Reactor

A laboratory-scale sequencing batch reactor (SBR) was used in this study. The reactor was originally inoculated with activated sludge obtained from the Nine Springs wastewater treatment plant in Madison, WI, which uses a variation of the University of Cape Town (UCT) process designed to achieve biological P removal without nitrate removal (20) and operates with high aeration rates (21). Details of the lab-scale operation under cyclic anaerobic and micro-aerobic conditions are provided in reference (8). Briefly, the SBR had a 2-liter working volume and was fed with synthetic wastewater containing acetate (500 mgCOD/L) as the sole carbon source (C:P molar ratio of 20). The synthetic wastewater was dispensed as two separate media; Media A contained the acetate and phosphate, whereas Media B supplied the ammonia (8). The reactor was operated under alternating anaerobic and low oxygen 8-h cycles. Each cycle consisted of 1.5 h anaerobic, 5.5 h micro-aerobic, 50 min settling and 10 min decanting. During the micro-aerobic stage, an on/off control system was used to limit the amount of oxygen pumped to the reactor (0.02 L/min) and to maintain micro-aerobic conditions in the mixed liquor (DO set point = 0.2 mg/L). The hydraulic retention time (HRT) and solids retention time (SRT) were 24 h and 80 days, respectively. The pH in the system was controlled to be between 7.0 and 7.5.

### Sample Collection and Analytical Tests

To monitor reactor performance, mixed liquor and effluent samples were collected, filtered through a membrane filter (0.45 μm; Whatman, Maidstone, UK) and analyzed for acetate, 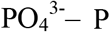, 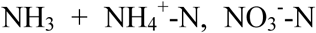, and 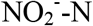. The concentrations of 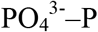 were determined according to Standard Methods (22). Total ammonia (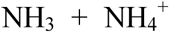) concentrations were analyzed using the salicylate method (Method 10031, Hach Company, Loveland, CO). Acetate, nitrite and nitrate were measured using high-pressure liquid chromatography as previously described (8).

For 16S rRNA-based tag sequencing and metagenomic analyses, biomass samples from the reactors were collected weekly and stored at −80°C until DNA extraction was performed. DNA was extracted using UltraClean^®^ Soil DNA Isolation Kit (MoBIO Laboratories, Carlsbad, CA). Extracted DNA was quantified using a NanoDrop spectrophotometer (Thermo Fisher Scientific, Waltham, MA) and stored at −80°C.

For transcriptomic analyses, biomass samples were collected across a single reactor cycle to capture key transition points in the EBPR cycle (Fig. 1). Samples (2 ml) were collected in microcentrifuge tubes, centrifuged, supernatant removed and cell pellets flash frozen in dry ice and ethanol bath within 3 min of collection. RNA was extracted from the samples using a RNeasy kit (Qiagen, Valencia, CA, USA) with a DNase digestion step. RNA integrity and DNA contamination were assessed using the Agilent 2100 Bioanalyzer (Agilent Technologies, Palo Alto, CA, USA).

**Figure 1.**
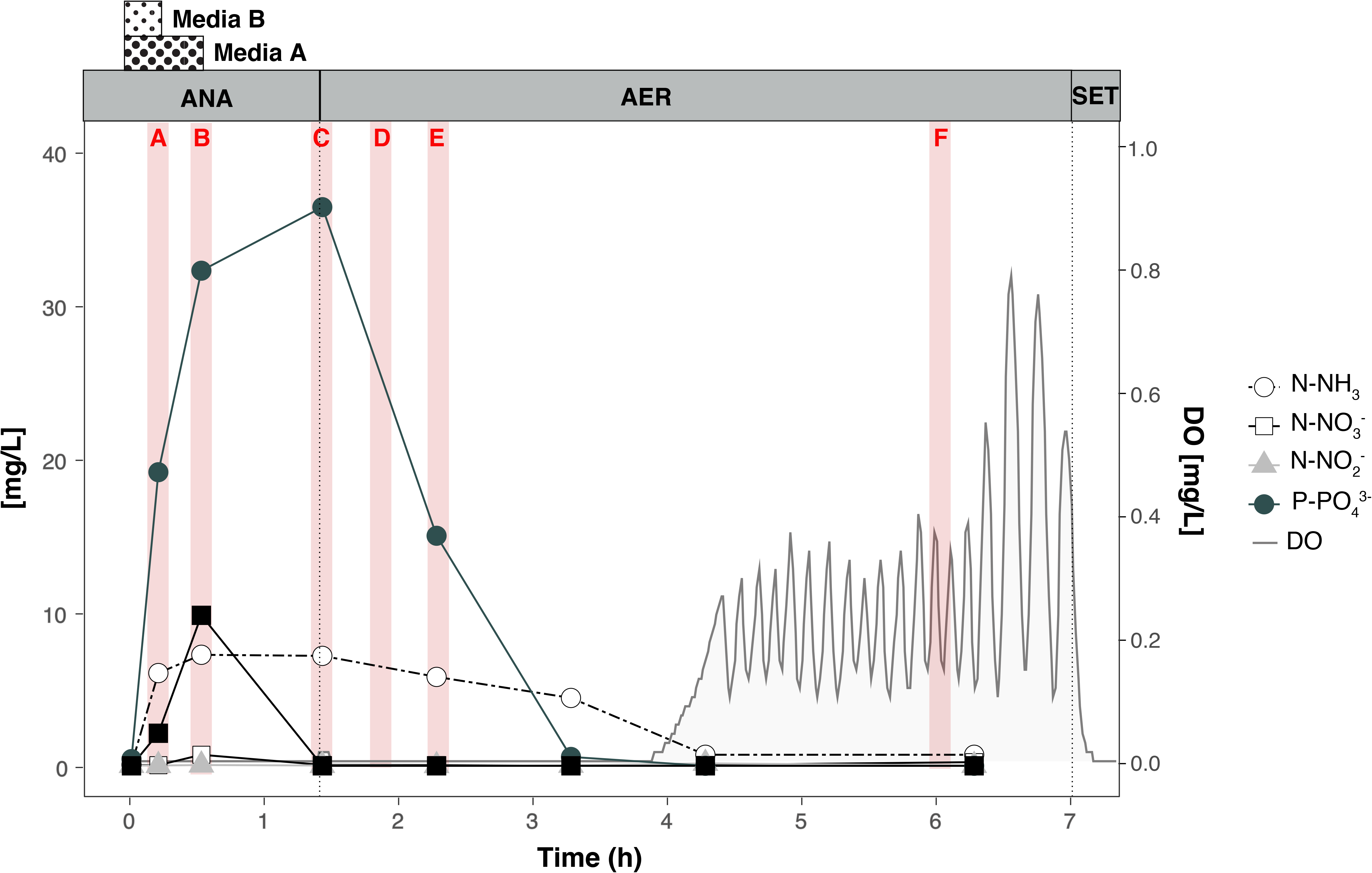
Nutrient profile of phosphorus, acetate, nitrogenous compounds and oxygen concentration in the lab-scale SBR on day-522. Dotted lines separate anaerobic (ANA), micro-aerobic (AER) and settling (SET) periods. Red bars indicate time-points used for RNA-seq (letters in red corresponds to samples names). The period of Media A (acetate and phosphate containing) and Media B (ammonia containing) addition are indicated for the anaerobic stage.

### Ribosomal RNA gene-based Tag sequencing

The composition of the microbial community in the reactor was determined via the analysis of high-throughput sequencing of 16S rRNA gene fragments. The hyper-variable V3–V4 regions of the bacterial 16S rRNA gene were amplified using the primers 515f/806r (23) as described in reference (8). The sequencing data is available under BioProject PRJNA482250. Briefly, PCR products were generated using the Ex Taq kit (Takara); cycling conditions involved an initial 5 min denaturing step at 95°C, followed by 35 cycles at 95°C for 45 s, 50°C for 60 s, 72°C for 90 s, and a final elongation step at 72°C for 10 min. Amplicons were visualized on an agarose gel to confirm product sizes. Purified amplicons were pooled in equimolar quantities and sequenced on an Illumina Miseq benchtop sequencer using pair-end 250 bp kits at the Cincinnati Children’s Hospital DNA Core facility.

Paired-end reads obtained were merged, aligned, filtered and binned into operational taxonomic units (OTU) with 97% identity using the QIIME pipeline (24). Chimeric sequences were removed using UCHIME (25). The most representative sequences from each OTU were taxonomically classified using the MIDas-DK database (26).

### Quantitative polymerase chain reaction (qPCR)

Quantification of each Accumulibacter clade was carried out by qPCR using a set of clade-specific primers targeting the polyphosphate kinase (*ppk1*) gene (8). All qPCR reactions were run in a LightCycler 480 system (Roche applied Science, Indianapolis, IN). Each reaction volume was 20 μL and contained 10 μL iQ^™^ SYBR^®^ Green Supermix (BioRAD Laboratories, Hercules, CA), 0.8 μL each of 10 μM forward and reverse primer, 4.4 μL nuclease free water and 4 μL of sample. Templates for qPCR were obtained from clone collections or gene synthesis (IDT, USA). In all cases, ten-fold serial dilutions of each template (ranging from 10^1^ to 10^7^ copies per reaction) were used to generate qPCR calibration curves. All samples were processed in triplicate and each reaction plate contained non-template controls.

### Metagenome sequencing, assembly and binning

Samples from days 522 and 784 were selected for metagenomic analysis. Illumina TruSeq DNA PCR free libraries were prepared for DNA extracts according to the manufacturer’s protocol and paired-end sequenced on either the Illumina HiSeq 2000 platform (v4 chemistry, 2 × 150□bp; 522-day sample), or the Illumina MiSeq platform (v3 chemistry, 2 × 250□bp; 784-day sample). A sample from day 522 was also sequenced on MinION (Oxford Nanopore Technologies, Oxford, UK), according to the Oxford Nanopore Genomic DNA Sequencing protocol (SQK-MAP003). The MinION flowcell was run for 48-h using the MinION control software, MinKNOW (version 47.3) and online base-calling was performed by the software Metrichor (version 2.23). Raw reads have been submitted to NCBI and are accessible under the BioProject identifier PRJNA322674.

Illumina unmerged reads were quality-trimmed and filtered with Sickle (https://github.com/ucdavis-bioinformatics/sickle.git), using a minimum phred score of 20 and a minimum length of 50 bp. Metagenomic reads from day 522 were assembled using the metaSPAdes pipeline of SPAdes 3.9.0 (27) and individual genome bins were extracted from the metagenome assembly using MaxBin (28). Genome completeness and redundancy was estimated using CHECKM 0.7.1 (29). Taxonomic identity of the bins was assigned using PhyloSift v 1.0.1 (30) and the script ‘parse_phylosift_sts.py’ available at https://github.com/sstevens2/sstevens_pubscrip/blob/master/parse_phylosift_sts.py (with options *-co_prob* 0.7 and *-co_perc* 0.7). Bin information is summarized in Table S1.

Two putative Accumulibacter bins (bin.046 and bin.097.4 in Table S1) were identified. The bin with the highest completeness (bin.046; hereafter referred to as UW-LDO-IC) was selected for further analysis and subjected to further processing to improve its quality. Redundant scaffolds were manually removed based on tetra-nucleotide frequency and differential coverage, using metagenomic reads from days 522 and 784 and following the anvi’o workflow described in A. M. Eren et al. (31). Further scaffolding was performed on UW-LDO-IC using Nanopore long-reads and the LINKS algorithm (32). Gapcloser (https://sourceforge.net/projects/soapdenovo2/files/GapCloser/) was used for additional gap filling. Table S2 displays quality metrics of the Accumulibacter draft genome after each of the steps previously described. The metagenomic assembly and final Accumulibacter bin was annotated using MetaPathways v 2.0 (33) and can be found under the GenBank accession number QPGA00000000.

### Average Nucleotide Identity (ANI)

Pair-wise ANI values of Accumulibacter genomes were obtained using the ANIm method (34) and implemented in the Python script ‘calculate_ani.py’ available at https://github.com/ctSkennerton/scriptShed/blob/master/calculate_ani.py

### Phylogenetic Analyses

The phylogeny of Accumulibacter UW-LDO-IC was assessed by constructing a phylogenetic tree using a concatenated alignment of marker genes. Published Accumulibacter draft and complete genomes were included in the analysis. First, PhyloSift was used to extract a set of 38 marker genes from each genome. Then, the extracted marker protein sequences were concatenated into a continuous alignment to construct a maximum-likelihood (ML) tree, using RAxML v 7.2.8 (35). RAxML generated 100 rapid bootstrap replicates followed by a search for the best-scoring ML tree.

For phylogenetic analyses of the polyphosphate kinase 1 (*ppk1*), nitrate reductase alpha subunit (*narG*), nitrite reductase (*nirS*), nitric oxide reductase (*norZ*), and nitrous oxide reductase (*nosZ*) genes, nucleotide datasets were downloaded from the NCBI GenBank database (36). Alignments were performed using the ‘AlignSeqs’ command in the DECIPHER “R” package (37). Phylogenetic trees were calculated using neighbor-joining criterion with 1,000 bootstrap tests for every node and the trees were visualized with the assistance of FigTree v1.4 (http://tree.bio.ed.ac.uk).

### RNA sequencing, filtering and mapping

Six biomass samples from within a reactor cycle on operational day 522 (Fig. 1) were collected and immediately processed to determine transcript abundance. RNA was extracted from the samples using a RNeasy kit (Qiagen, Valencia, CA, USA) with a DNase digestion step. RNA integrity and DNA contamination were assessed using the Agilent 2100 Bioanalyzer (Agilent Technologies, Palo Alto, CA, USA). Ribosomal RNA (rRNA) was removed from 1 μg of total RNA using Ribo-Zero rRNA Removal Kit (Bacteria) (Epicentre, Madison, WI, USA). Libraries were generated using the Truseq Stranded mRNA sample preparation kit (Illumina, San Diego, CA, USA), according to the manufacturer’s protocol. The libraries were quantified using KAPA Biosystem’s next-generation sequencing library quantitative PCR kit and run on a Roche LightCycler 480 realtime PCR instrument. The quantified libraries were then prepared for sequencing on the Illumina HiSeq 2000 sequencing platform utilizing a TruSeq paired-end cluster kit, v3, and Illumina’s cBot instrument to generate a clustered flowcell for sequencing. Sequencing of the flowcell was performed on the Illumina HiSeq 2000 sequencer using a TruSeq SBS sequencing kit 200 cycles, v3, following a 2×150 indexed run recipe. Sequence data were deposited at IMG/M under Taxon Object IDs 3300004259-3300004260 and 3300004621-3300004624.

RNA reads were quality filtered and trimmed with Sickle and forward and reverse reads were merged using FLASH (v. 1.2.11) (38). Ribosomal RNA sequences were removed with SortMeRNA using six built in databases for bacterial, archaeal and eukaryotic small and large subunits (39). Reads that passed filtering were then mapped to the metagenomic assembly and to the complete and draft genomes using the BBMap suite (40) with default parameters, respectively. Read counts were then calculated using HTseq with the ‘intersection strict’ parameter (41). Read counts were normalized by total reads in the sequencing run, the number of reads that remained after rRNA filtering, and the fraction of total reads that aligned to the assembly and genomes (Table S3) in each sample. Number of reads mapping to each gene were then converted to log_2_ reads per kilobase per million (RPKM (42)).

### Primer design

PCR primer sets targeting the Accumulibacter UW-LDO-IC’s genes *nirS, narG, norZ, nosZ,* the *ccoN* subunit of *cbb3* and the *ctaD* subunit of *caa3* cytochrome oxidases, were designed to quantify expression of these genes in cDNA samples from the reactor. For comparison, primers for the *rpoN* gene were also designed (Table S4). For primer design, genes from UW-LDO-IC and other published Accumulibacter genomes were aligned with homologs from bacteria that share relatively high DNA sequence identity with Accumulibacter. The list of aligned gene sequences was then submitted to DECIPHER’s Design Primers web tool (43) using the following parameters: primers length ranging from 17-26 nucleotides with up to 2 permutations and PCR product amplicon length between 75-500 bp.

PCR amplification of UW-LDO-IC gene fragments was carried out on extracted genomic DNA from the lab-scale SBR, in a 25 μL reaction volume with 400 nM of each forward and reverse primer. The PCR program consisted of an initial 10-min denaturation step at 95°C, followed by 30 cycles of 95°C for 30 s, 64°C for 30 s, and 72°C for 30 s, and then a final extension at 72°C for 5 min. The presence and sizes of the amplification products were determined by agarose (2%) gel electrophoresis of the reaction product. The amplified fragments were then purified, cloned using a TOPO TA cloning kit (Invitrogen, CA) according to the manufacturer’s instructions. Fragments were single-pass Sanger sequenced, and the sequences were aligned to Accumulibacter UW-LDO-IC to confirm specificity. In all cases, PCR fragment sequences aligned to the corresponding gene in UW-LDO-IC with a percent of identity > 97%, whereas the identity percentage with other Accumulibacter genomes was < 95%. The sequences have been deposited in GenBank under BioProject PRJNA482254.

### Quantitative real-time PCR

Complementary DNA (cDNA) was generated from 500 ng of total RNA, primed by random hexamers (SuperScript II first-strand synthesis system, Invitrogen, Carlsbad, CA, USA). The reaction was terminated by incubation at 85°C for 5 min and RNase H treatment was performed to degrade RNA in RNA:DNA hybrids. Subsequently, 4 μL of 10x diluted cDNA was applied as the template in qPCR. All quantifications were performed in triplicate. The qPCR was conducted on a LightCycler 480 (Roche, Switzerland) using iQ SYBR Green Supermix (Bio-Rad) with a total reaction volume of 20 μL. All qPCR programs consisted of an initial 3 min denaturation at 95°C, followed by 45 cycles of denaturing at 95°C for 30 s, 64°C for 30 s, and 72°C for 30 s. RNA samples without reverse transcription were used as no RT (reverse transcription) controls to evaluate DNA contamination for all primers tested. The relative fold change of target gene expression between samples was quantified using Accumulibacter *rpoN* as a reference gene.

### Operons and upstream motif identification

*De novo* motif detection analysis was conducted on the intergenic regions upstream of the *nar, nir, nor, nos, caa3* and *cbb3* operons of Accumulibacter UW-LDO-IC using MEME (44). FNR-motif sites were further identified in both strands of other Accumulibacter genomes using the FIMO tool (45) (p-values < 1 x 10^-5^). Search was limited to the promoter region, represented by the 300 bp intergenic region upstream of the transcriptional start site, of all protein-coding sequences annotated with MetaPathways (33).

Putative operons were determined using the same set of criteria as in reference (46). That is, each operon enclosed adjacent genes with the same orientation, co-expressed with a minimum Pearce correlation coefficient of 0.7, and an intergenic region between genes of 1000 base pairs or less.

## RESULTS AND DISCUSSION

### Characterization of reactor operation and Accumulibacter community structure

A nutrient profile of one reactor’s cycle from the date samples were collected for transcriptomics (day 522) is shown in Fig. 1. Acetate was slowly added to the reactor during the first 32 minutes of the anaerobic phase, with no accumulation observed, as it was rapidly consumed. P release to the mixed-liquor was observed during the period of acetate uptake. During the anaerobic stage, ammonia-containing media was supplied during the first 16 min of the anaerobic stage and the ammonia accumulated in the reactor. In the micro-aerobic phase, when measured DO was about 0.02 mg/L, P was taken up by cells. Simultaneously, nitrification occurred during the first 3 hours of aeration, without 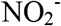 or 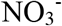 accumulation, indicating simultaneous nitrification and denitrification in the reactor. After all the substrates that exert oxygen demand were depleted, oxygen increased and fluctuated around the 0.2 mg/L set point. These observations are consistent with an efficient EBPR process under cyclic anaerobic and micro-aerobic conditions, as discussed elsewhere (8).

Samples collected on the same day for 16S rRNA amplicon sequencing indicated that Accumulibacter was the most abundant bacterium in the reactor, accounting for 34% of the total number of reads (Table S5). Members of the *Competibacteraceae* family (16%) and the *Lewinella* genus (11%) were also abundant. The diversity within the Accumulibacter lineage was assessed by qPCR (Table S6). In these samples the Accumulibacter members were dominated by Clade IC, which accounted for 74% of the total, followed by Clade IID (14%) and IIA (9%). As described before (8), clade IC predominated in the reactor during at least 300 days of operation, with abundances greater than 87% of total Accumulibacter, and batch tests suggested its ability to use oxygen, nitrite, and nitrate as electron acceptors (8). Clade IC has also been described as the dominant Accumulibacter clade in a reactor operated under anaerobic/anoxic/oxic conditions (18). However, contrary to our findings, batch tests suggested that this strain was not capable of using nitrate as external electron acceptor for anoxic P-removal. This inconsistency among denitrifying capabilities could be the result of genetic variations that are not captured with the current *ppk1*-based clade definitions.

### Assembling a draft genome of Accumulibacter clade IC

All existing metagenome assembled genomes (MAGs) of Accumulibacter have been obtained from bioreactors operated under conventional anaerobic/aerobic cycles that use abundant aeration (47-50). To date, only one genome of Accumulibacter has been closed (clade IIA strain UW-1), while draft genomes from 5 different clades (clade IA, IB, IIA, IIC and IIF) have been reconstructed from metagenomic data. Since the SBR reactor operated with anaerobic/micro-aerobic cycles enriched for a less common clade of Accumulibacter, we performed a metagenomic assessment of the microbial community in the reactor. The whole-community DNA of two samples from the reactor were sequenced using two different technologies: Illumina and Oxford Nanopore. Short Illumina reads were initially assembled and binned into 136 different bacterial draft genomes (Table S1). One of these bins was classified as Accumulibacter (bin.046) and characterized by its high completeness (94.8%) and relatively high redundancy (28.9%), likely due to the presence of redundant gene markers from other Accumulibacter strains. The presence of another incomplete bin also classified as Accumulibacter (bin.097.4; 26.0% completeness) further support the idea of other Accumulibacter strains present at lower concentrations, in agreement with the assessment of diversity based on the *ppk1* gene (Table S6). To obtain a higher-quality draft genome of the dominant Accumulibacter strain, the differential coverage of two metagenomic samples was used to remove contaminant contigs, reducing the redundancy level to 0.84% of marker genes. Finally, Nanopore sequencing data was used for further scaffolding; gaps were filled using the GapCloser tool. The resulting near-complete draft genome, termed Accumulibacter sp. UW-LDO-IC, has 4.7 Mbp in total with average GC content of 62.5% (Table S2) and encoded 95.2% of marker genes with 0.68% redundancy.

A phylogenetic tree constructed from the *ppk1* gene encoded in UW-LDO-IC, other Accumulibacter genomes and sequences available at NCBI, were used to classify UW-LDO-IC into one of the 14 Accumulibacter clades described to date. According to the phylogenetic tree topology of the *ppk1* gene (Fig. 2A), UW-LDO-IC’s *ppk1* clustered with sequences previously classified as clade IC, and therefore, the draft genome assembled herein would belong to this clade. With more Accumulibacter draft genomes becoming available in recent years, the tree topology also suggests that Accumulibacter BA-92 (49), a draft genome initially classified as clade IC may be better classified as belonging to clade IB along with the draft genome HKU-1 (48). To further evaluate this potential misclassification, a phylogenetic tree of a concatenated protein alignment of 38 universally distributed single-copy marker genes (51) was constructed (Fig. 2B). This tree topology supports the classification of UW-LDO-IC as belonging to Type I, but separate from Accumulibacter BA-92 and HKU-1. Thus, we propose that UW-LDO-IC be classified as the only draft genome representing clade IC, and Accumulibacter BA-92 and HKU-1 be classified in clade IB, along with Accumulibacter UBA2783 (52).

**Figure 2.**
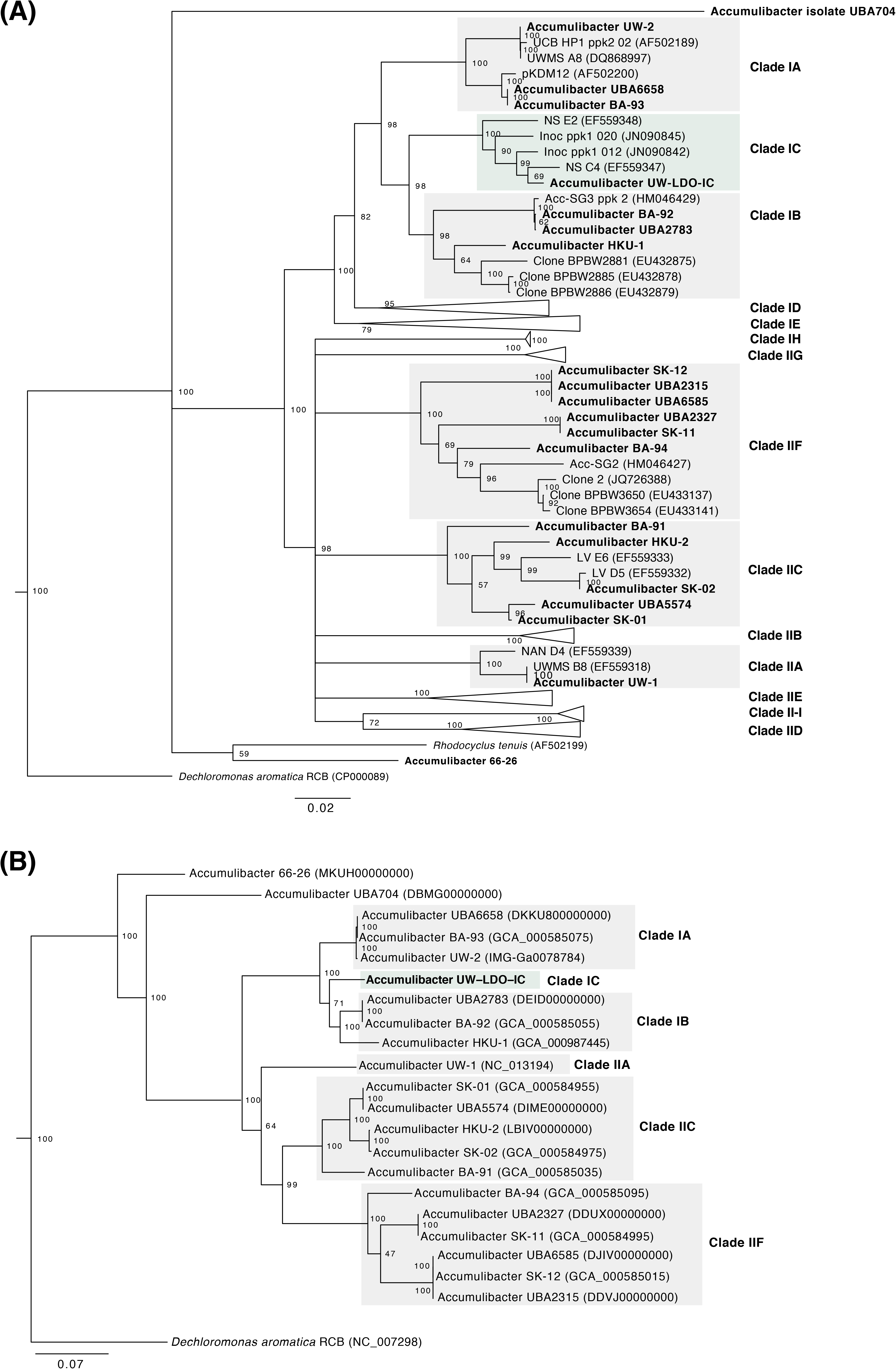
Phylogeny of Accumulibacter UW-LDO-IC. (A) Neighbor-joining phylogenetic tree based on nucleic acid sequences of *ppk1* found in Accumulibacter genomes. (B) RAxML phylogenetic tree of a concatenated alignment of 38 marker genes (nucleotide sequences) of the Accumulibacter genus. Bootstrap values are shown in the tree branches based on 1000 and 100 bootstrap replicates, respectively. The scale bar represents the number of nucleotide substitutions per site.

Average nucleotide sequence identity (ANI) between UW-LDO-IC and formerly published Accumulibacter genomes was used to confirm the phylogenetic analysis, as this method has been shown to correlate well with previously defined species boundaries (53, 54). The calculated ANI and alignment fraction for the Accumulibacter genomes showed that UW-LDO-IC shares only 88.7 and 88.3% identity and 67.2 and 60.2% alignment with Accumulibacter HKU-1 (Clade IB) and BA-92 (Clade IB), respectively (Fig. S1). The low ANI and low alignment, as well as the concatenated markers phylogeny, indicates that Accumulibacter UW-LDO-IC has significant differences with other Accumulibacter genomes, none of which have been retrieved from BNR microbiomes adapted to minimal aeration.

### Denitrification potential of Accumulibacter UW-LDO-IC

A comparison of the genetic inventory involved in anoxic respiration revealed differences between UW-LDO-IC and previously published Accumulibacter genomes (Table 1 and S7). Among the differences found is that UW-LDO-IC encodes a full denitrification pathway, which involves a membrane bound nitrate reductase of the NarG type (*narGHJI* operon), as well as a nitrate/nitrite transporter homologous to the *narK* gene, a periplasmic cytochrome *cd_1_* nitrite reductase NirS and the proteins involved in heme d_1_ biosynthesis (*nirMCFDLGHjN*), a quinol-dependent nitric oxide reductase *norZ* and a nitrous oxide reductase Nos (*nosZDFYL*) (Table 1). The genetic context of these genes was compared to other Accumulibacter genomes (Fig. S2). The position of denitrifying genes within the genome varied among different clades. Overall, genes flanking the *nar* operon, *nirS-2* and *nosZ* genes were the same in all Accumulibacter clades where these genes were identified, including UW-LDO-IC. Unlike clade IIC or IB, *nirS-1* of UW-LDO-IC was not positioned next to the *nar* or *nor* genes, but the genomic context of *nirS-1* in UW-LDO-IC differed from any Accumulibacter genome. Since *nirS-1* from UW-LDO-IC presented high identity with Accumulibacter HKU-1 and BA-92 (91%), this difference in the genome context might be caused by lateral gene transfer. The presence of transposase genes next to *uspA* in BA-93 (a flanking gene of *nirS-1* in UW-LDO-IC (Fig. S2)) supports this hypothesis.

**Table 1.**
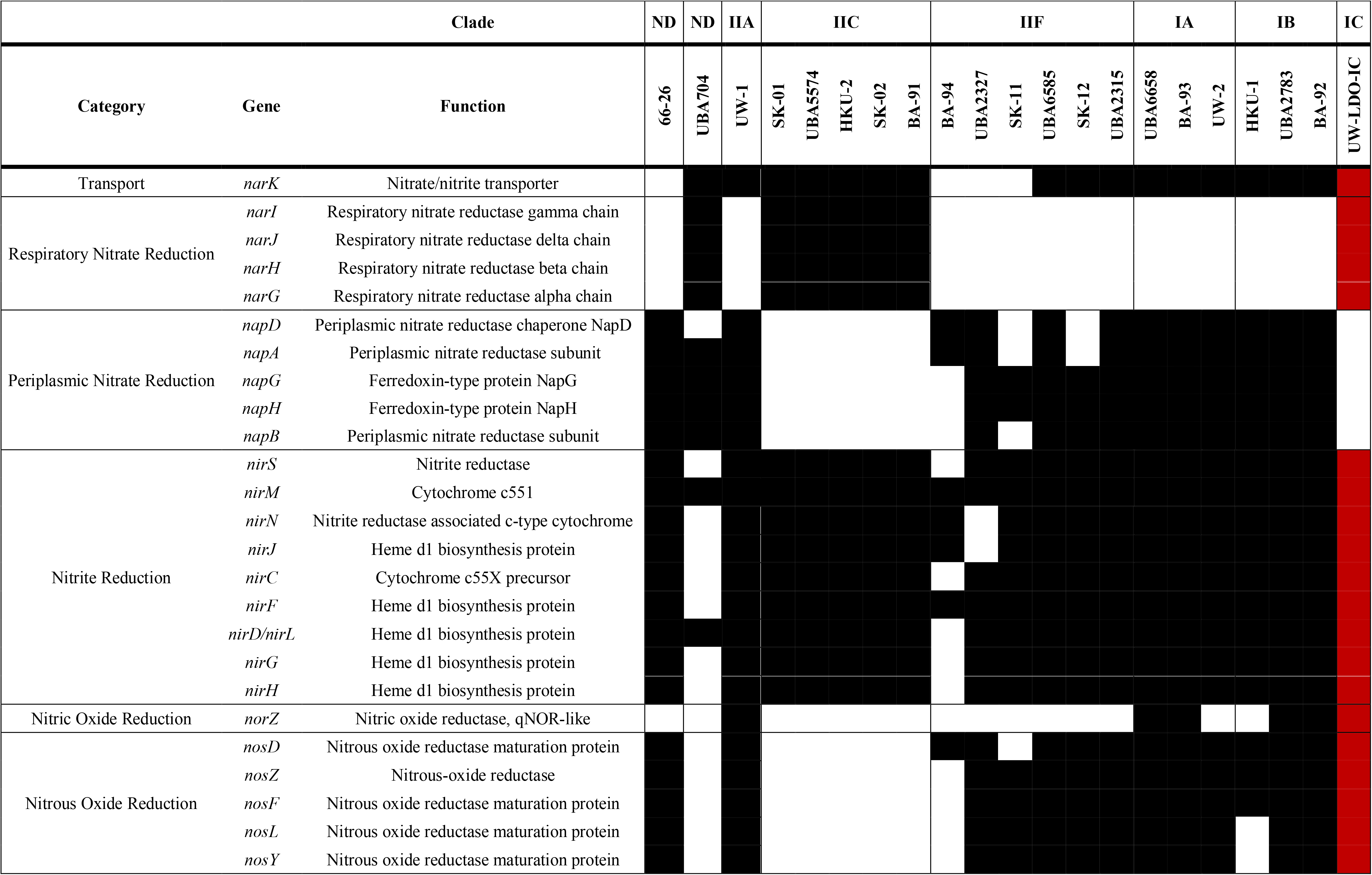
Inventory of genes associated with denitrification in complete and draft genomes of *Accumulibacter*. Black and white rectangles represent presence and absence of each gene, respectively. Genes present in the assembled Accumulibacter sp. UW-LDO-IC genome are highlighted in red. ND: Not determined.

An alignment of full-length sequences of subunits *narG, nirS, norZ* and *nosZ,* which contain the active sites in the corresponding enzymes, was used to evaluate the phylogenetic associations of these genes (Fig. S3-S6). This analysis revealed that all UW-LDO-IC genes involved in denitrification were closely related to other members of the Accumulibacter genus, ruling out possible contig contamination from other denitrifying bacteria during binning. Phylogenetic differentiation between genes belonging to type I and II of Accumulibacter was observed, where genes belonging to UW-LDO-IC clustered with other genes from type I genomes. In the case of *narG*, which has only been identified in genomes from clade IIC, UW-LDO-IC clustered separatedly from the Clade II genes (Fig. S3).

Denitrifying genes encoded in Accumulibacter genomes were phylogenetically related to different taxonomic groups. Genes encoding the NarG proteins seem to have derived from other *Betaproteobacteria*. Interestingly, these genes exhibit phylogenetic relation with *narG* from different bacterial families (*Comamonadaceae, Pseudomonaceae, Burkholderaceae and Rhodocyclaceae*), indicating a similar origin (Fig. S3). The close relationship of Accumulibacter’s *narG* with the plasmid-encoded gene in *Burkholderia phymatum* suggests potential mobility of this gene across genera.

Multiple copies of *nirS* are present in the majority of Accumulibacter genomes, including UW-LDO-IC (Fig. S4). The phylogenetic analysis of sequences encoding this gene indicates two main clusters of *nirS* including sequences from both Accumulibacter and other members of the *Rhodocylaceae* family, with one of these clusters, harboring the *nirS-2* and *nirS-3* genes of UW-LDO-IC, being closely related to sequences from the *Pseudomonas* genus. Interestingly, no other *Rhodocyclaceae* member encoded a quinol-dependent nitric oxide reductase (*norZ*), since in these species, nitric oxide reduction requires the activity of a cytochrome bc-type complex (*norBC*). Accumulibacter’s *norZ* was instead phylogenetically most closely related to *Polaromonas*, another member of the *Comamonadaceae* family (Fig. S5). Finally, the closest sequences to Accumulibacter’s *nosZ* gene, belonged to the *Dechloromonas* (*Rhodocyclaceae* member) (Fig. S6). These findings are in agreement with a recent ancestral genome reconstruction and evolutionary analysis of the Accumulibacter lineage, which found that the denitrification machinery was not present in the last common ancestor of Accumulibacter and that one of the most abundant source of horizontally transferred genes are the *Burkholderiales*, including many from *Comamonadaceae* (see Supplementary Table 6 in reference (55)). Overall, these results suggest that part of the denitrification machinery of Accumulibacter was laterally transferred from other microorganisms commonly found in activated sludge.

Only incomplete denitrification pathways, with the potential of reducing nitrite to nitrogen gas (Accumulibacter UW-1, UBA6658, UBA2783, UW-2, BA-92 and BA-93) and nitrate to nitric oxide (Accumulibacter SK-01, SK-02, BA-91, UBA5574 and HKU-2), were identified in other Accumulibacter genomes (Table 1). Furthermore, evidence for a periplasmic nitrate reductase Nap enzyme was not found in UW-LDO-IC but is present in other Accumulibacter genomes (Table 1). Although it has been hypothesized that Nap could be responsible for the nitrate reduction step in some Accumulibacter strains (49), due to the functional diversity of this enzyme, involved in denitrification, nitrate reduction to ammonia, maintenance of cellular oxidation-reduction potential and nitrate scavenging, the presence of a *nap* homolog in an organism’s genome cannot necessarily be linked to nitrate respiration (56). For instance, despite the existence of a *nap* operon within the Accumulibacter UW-IA (clade IIA) genome (Table 1 and S7), J. J. Flowers et al. (10) demonstrated through batch denitrification assays, that this strain could not use nitrate as a terminal electron acceptor. Therefore, UW-LDO-IC would be the first Accumulibacter genome found to encode a full denitrifying machinery, containing genes directly involved in the reduction of nitrate to nitrogen gas, settling a long-running debate in the research community about whether Accumulibacter can achieve complete nitrate reduction to nitrogen gas while cycling polyphosphate (10, 15, 17, 57).

### Aerobic respiration potential of Accumulibacter UW-LDO-IC

The presence of known terminal oxidases, which catalyze oxygen reduction to water during the final step of the electron transport chain (58), was examined in the available Accumulibacter genomes. Three known terminal oxidases were annotated in all genomes of Accumulibacter: cytochrome *aa_3_* (encoded by the *ctaDCE*), *ba_3_* (encoded by the subunits *cbaA and cbaB*) and *cbb_3_* (encoded by operon *ccoNOQP*) oxidases, which accept electrons from cytochrome c and transfer them in reactions involved in oxygen reduction (Table 2 and S8). The *aa_3_*-type oxidases have low affinity for oxygen and usually play a dominant role under high oxygen conditions (59-61). Phylogeny of the first subunit of this enzyme (Fig. S7) shows marked differentiation of the Accumulibacter lineage from other *Rhodocyclaceae* organisms, although some genomes of Accumulibacter, including UW-I (clade IIA), clustered closely to *Dechloromonas*. All genomes classified as clade IIF lacked this subunit (Table 2), and thus, members of this group might rely on other cytochrome c oxidases for aerobic respiration.

**Table 2.**
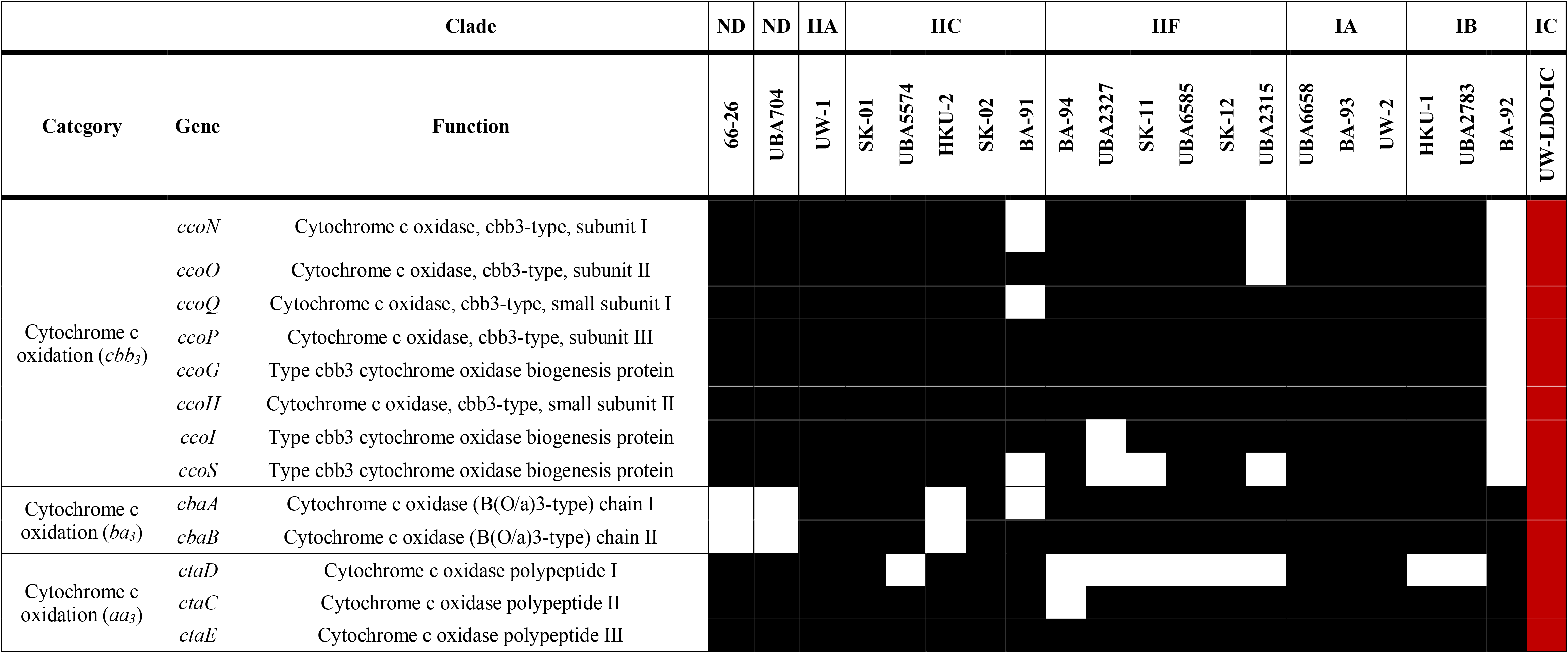
Inventory of genes associated with aerobic respiration in complete and draft genomes of Accumulibacter. Black and white rectangles represent presence and absence of each gene, respectively. Genes present in the assembled Accumulibacter sp. UW-LDO-IC genome are highlighted in red. ND: Not determined

On the other hand, *cbb*_3_, oxidases are known to have very high affinity for oxygen and to be induced under low oxygen conditions in many bacteria (62-66). This enzyme is widespread in the Accumulibacter lineage, since 18 out of the 21 genomes analyzed herein harbored the *cco* operon encoding the subunits of this enzyme (Table 2). These results indicate that the ability to survive in low-oxygen concentrations is common across Accumulibacter strains and would explain why lowering oxygen concentration does not seem to negatively affect EBPR (8, 21, 67). The phylogenetic analysis of the first subunit of this enzyme, *ccoN*, shows conservation of this trait among members of the *Rhodocyclaceae* family (Fig. S8).

Lastly, *ba_3_*-type cytochrome oxidase has been mostly studied in the extremophile bacterium *Thermus thermophilus*, where it is usually expressed under oxygen limiting conditions (68), but little is known about its role in other organisms. In Accumulibacter this enzyme is present in all clades (Table 2) and its phylogeny reveals that it could be derived from non-*Rhodocyclaceae* members (Fig. S9).

### Changes in transcript abundance during an anaerobic/micro-aerobic cycle

Transcriptional investigations in Accumulibacter have illuminated how the complex and unique metabolism of this lineage is a result of highly dynamic gene expression (46, 48, 69, 70). Recently, the power of time series metatranscriptomics was used to analyze gene expression patterns in Accumulibacter during an anaerobic-aerobic EBPR cycle (46). This study was carried out with high oxygen concentrations in a reactor where nitrification was inhibited and anoxic respiration did not take place. In order to study the effect of limited oxygen conditions in the expression of the respiratory machinery of Accumulibacter UW-LDO-IC, we used a similar time series high-resolution RNA-seq approach and contrasted the effect of high/low oxygen in the metabolism of this strain.

A time-series metatranscriptomic dataset of the lab-scale reactor was obtained in collaboration with DOE-JGI. Samples were collected at the beginning of the anaerobic stage and at different times during the micro-aerobic stage when DO conditions were ~0.05 mg/L and 0.25 mg/L, respectively (Fig. 1). RNA sequencing resulted in 1,718,478,214 total reads across the six samples (Table S3). Quality filtering, merging, and rRNA removal resulted in 396,995,401 sequences for downstream analysis. Resulting reads were then competitively mapped to Accumulibacter UW-LDO-IC and other publicly available Accumulibacter complete and draft genomes, including clades IA, IB, IIA, IIC and IIF (Table S9). Between 48-50% of each sample’s filtered RNA reads mapped to the genome of UW-LDO-IC, indicating that this was the most active bacterium in the community. No other genome of Accumulibacter retrieved more than 0.52% of mapping reads, and therefore, strains closely related to other available Accumulibacter genomes were not active members of the community.

Transcripts mapping to genes related to denitrification were investigated by analyzing Accumulibacter UW-LDO-IC gene expression patterns during the cycle. Fig. 3 depicts the relative expression of genes encoding the *nar*, *nir*, *nor* and *nos* operons, and nitrite-nitrate transporters (*narK*) at each time point; the minimum expression of each gene was subtracted from each point to allow better visualization of the dynamics of each gene over the course of the cycle. All subunits of the *narGHJI* operon showed similar patterns, with transcript abundance increasing during the anaerobic stage, followed by a decrease in transcript levels as the oxygen concentration increased in the system at the end of the cycle (Fig. 3A). Only one of the three *nirS* copies (*nirS*-1) present in Accumulibacter UW-LDO-IC showed an increment on its transcript abundance during the cycle (maximum Δlog_2_(RPKM) read count >1), with a pattern similar to that of the *narGHJI operon* (Fig. 3B). Transcripts from the *narK* and *nosZ* genes also increased during the anaerobic stage, but their abundance started decreasing as soon as air was introduced to the reactor (Fig. 3D-E). The gene *norZ* did not display a notable change in relative transcript abundance (maximum Δlog_2_(RPKM) read count < 1), although its expression increased over time (Fig. 3C). These observations are consistent with upregulation of denitrifying genes during the anaerobic stage, suggesting that oxygen concentration plays an important role in transcriptional regulation of these genes, as has been previously described (71).

**Figure 3.**
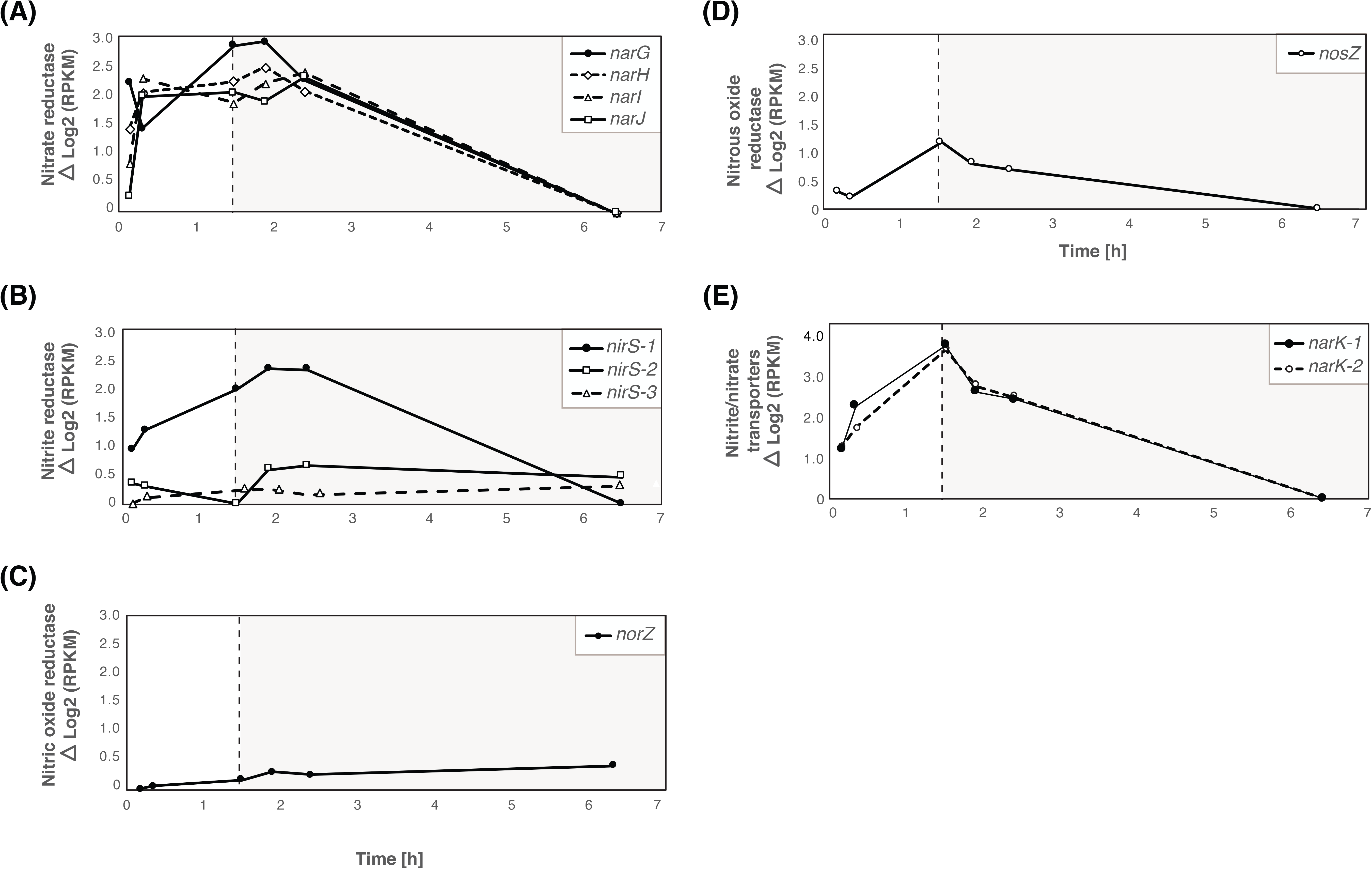
Gene expression profile patterns of denitrifying-related genes. Relative transcript abundances of (A) nitrate, (B) nitrite, (C) nitric oxide and (D) nitrous oxide reductase genes, and (E) nitrite/nitrate transporters in Accumulibacter UW-LDO-IC during the anaerobic (white panel) and micro-aerobic (grey panel) phases. Each time point’s expression value was normalized to the minimum expression of each gene over the cycle.

Similar transcript trends for denitrification-associated genes have been previously described in Accumulibacter, in reactors operating with anaerobic/aerobic cycles that used high aeration rates and where nitrification was chemically inhibited with allylthiourea (46, 69). In general, these studies showed upregulation of denitrification genes during anaerobic conditions and a reduction in transcript abundance when oxygen was introduced to the system. However, unlike the results observed at high-DO (46), our observations indicate that transcript abundance remains high after oxygen addition, when simultaneous nitrification/denitrification is occurring. The *norZ* gene expression pattern under low-oxygen also differs from the one reported at high-DO (46), since the transcripts of this gene in UW-LDO-IC did not considerably change during the aerobic period, whereas transcripts in UW-1 exhibited high variations during the operational cycle (Fig, 4B). Furthermore *nosZ* transcript levels under oxygen-limited conditions displayed a slower decrement rate than what was reported at high-DO (46), where negligible expression was observed after 1 hour of aeration, and similar results were described in S. M. He and K. D. McMahon (69) (Fig, 4C). Since the complete denitrification pathway was not present in the Accumulibacter UW-1 genome, no information about *nar* operon expression was available prior to our study.

**Figure 4.**
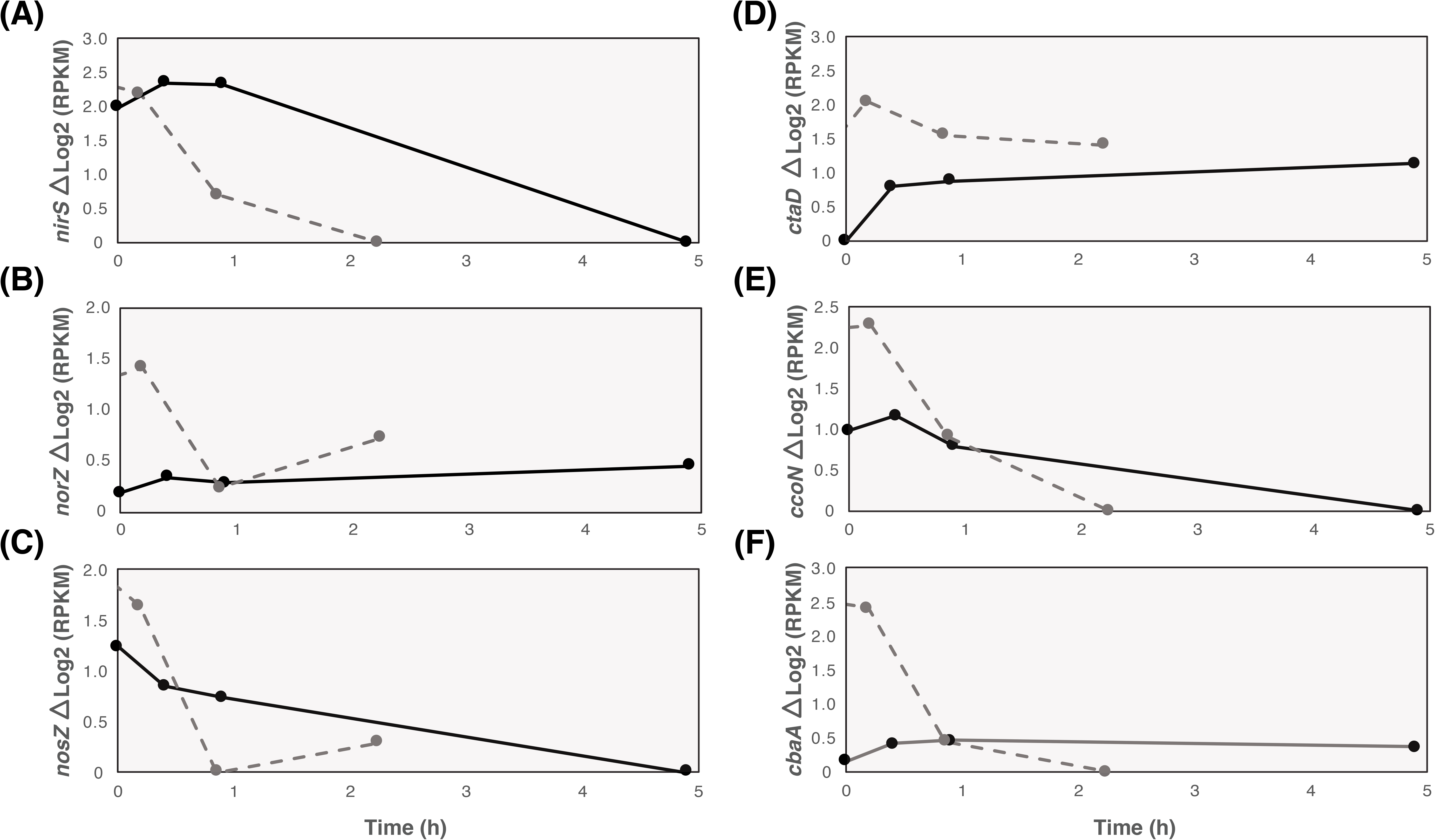
Gene expression comparison among Clade IC and IIA. Normalized transcript abundance of denitrification-related genes: (A) *nirS*, (B) *norZ* and (C) *nosZ*; and aerobic respiration-related genes: (D) *aa_3_* cytochrome subunit I, (E) *cbb_3_* subunit ccoN and (F) *ba_3_* subunit *cbaA* of Accumulibacter UW-TNR-IC (solid line) and UW-1 (dotted line) (46) during the aerobic stage of an EBPR cycle.

Previous studies have confirmed Accumulibacter’s capability to synthesize denitrification-associated proteins. In J. J. Barr et al. (47), nitrite and nitrous oxide reductase enzymes were detected by metaproteomic analysis of an Accumulibacter-enriched microbial community. No nitric oxide reductase protein was detected in this study, despite the presence of the gene *norZ i*n the genome of the strain enriched in this system, Accumulibacter BA-93 (IA). As observed in Fig. 3C, *norZ* transcript levels in UW-LDO-IC did not considerably changed over time, potentially indicating a low transcriptional level response to changes in oxygen or nitrite/nitrate concentrations, hence other regulatory mechanisms might control synthesis of this enzyme. Further experiments are still needed to understand whether post-transcriptional regulation would be also controlling denitrification-associated protein synthesis in Accumulibacter.

Changes in the expression of terminal oxidases in UW-LDO-IC were also identified. All subunits of the terminal cytochrome c oxidase *aa_3_* were upregulated during the entire micro-aerobic phase (Fig. 5A), indicating active aerobic respiration by this microorganism. Transcriptomic results also showed that the *cbb_3_*-type cytochrome oxidase transcripts decreased when DO increased in the reactor (Fig. 5B). Changes in cytochrome *ba_3_* oxidase transcript levels were less pronounced than those observed in the other terminal oxidases (maximum Δlog_2_(RPKM) read count < 0.5) (Fig. 5C), likely indicating a minor role of this enzyme during redox condition variations.

**Figure 5.**
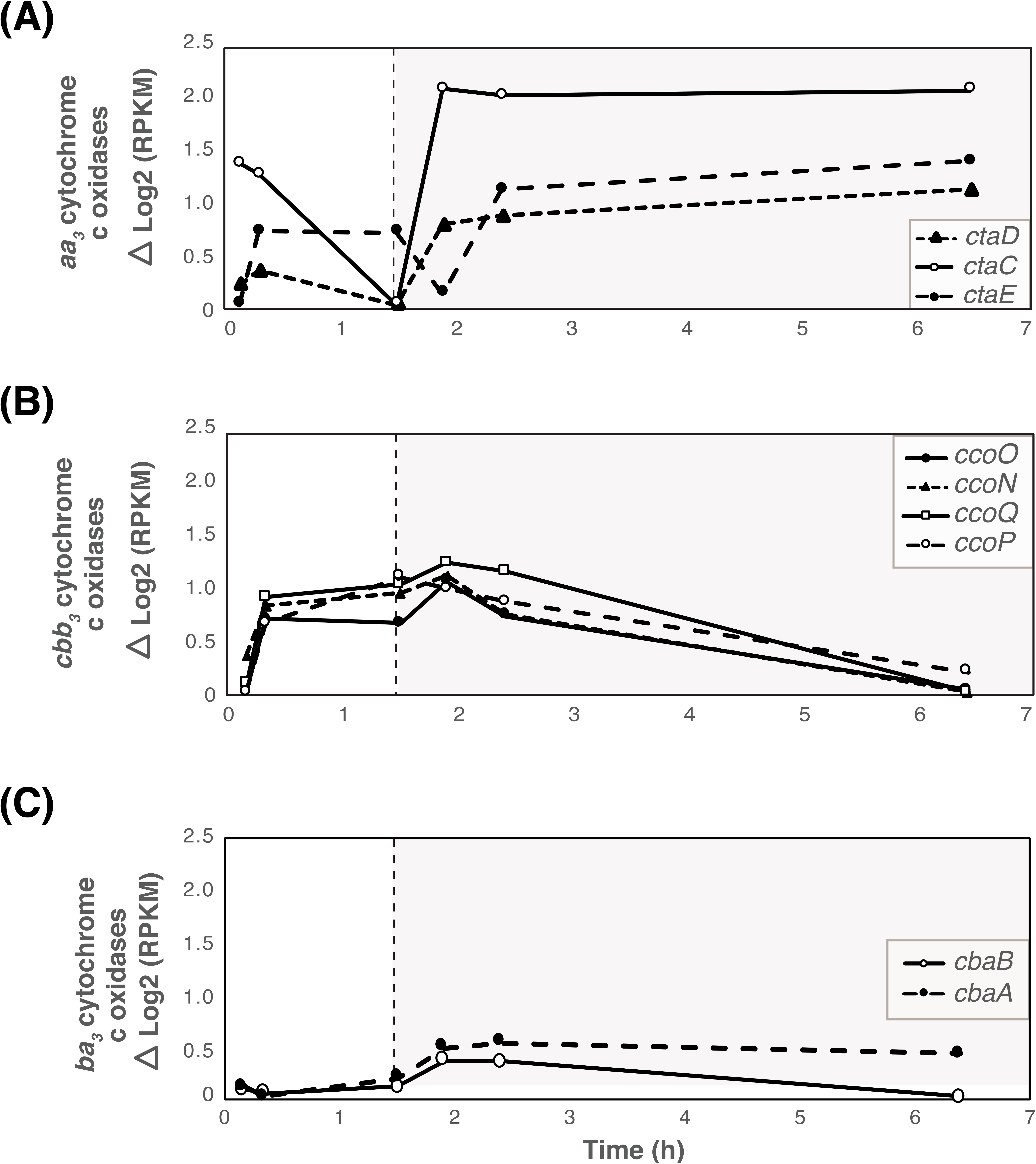
Gene expression profile patterns of aerobic respiration-related genes. Relative transcripts abundance of (A) *aa_3_*, (B) *cbb_3_* and (C) *ba_3_* cytochrome c oxidases in Accumulibacter UW-TNR-IC during the anaerobic (white panel) and micro-aerobic (grey panel) phases. Each time point’s expression value was normalized to the minimum expression of each gene over the cycle.

We compared the expression of these genes during high and low oxygen concentrations using data from reference (46) (Fig. 4). In both cases, the transcriptional expression of the low-affinity *aa_3_* cytochrome oxidase increased during the aerobic stage and remained upregulated until the cycle end (Fig. 4D). On the other side, *cbb_3_*-type cytochrome oxidase transcript abundance drastically decreased after turning on aeration in the high-aerated system, whereas the same gene in UW-LDO-IC remained upregulated during the first hour of minimal aeration, corroborating its function as a terminal oxidase induced by limited-oxygen conditions (Fig. 4E). Unlike UW-LDO-IC, Accumulibacter UW-1 had high *ba_3_* cytochrome c oxidase expression during the anaerobic stages, which declined after aeration started (Fig. 4F). Differences in the expression profile of the latter cytochrome suggests differential gene expression control among these two clades.

Overall, these results demonstrated how expression patterns of the genes responsible for denitrification and aerobic respiration, specifically *nar*, *nir* and *cbb3*, showed upregulation during the beginning of the micro-aerobic stage, supporting our hypothesis that Accumulibacter UW-LDO-IC is a denitrifying microorganism capable of simultaneously respiring oxygen and nitrate under micro-aerobic conditions.

### Validation of RNA-seq with RT-qPCR

RT-qPCR was conducted on six genes related to anoxic and aerobic respiration (*narG, nirS-1, norZ, nosZ, ccoN and ctaD*) to validate RNA sequencing results (Fig. 6). The no-reverse transcription control (NRTC) was used to evaluate the background caused by trace DNA contamination. The average of the difference in Ct (threshold cycle) values between the cDNA and NRTC control was 10 cycles, indicating that DNA contamination was negligible. The copy number of the RNA polymerase sigma-54 factor, encoded by the *rpoN* gene, was used as a reference gene for normalization of the RT-qPCR data, since this gene showed no significant changes (ΔRPKM < 1) in the RNA sequencing results and has been previously used as a reference gene for qPCR normalization in other bacteria (72). The RT-qPCR transcriptomic profile in Fig. 6 was obtained by normalizing each point by the minimum number of copies across the cycle. In all cases, genes regulation trends identified by RT-qPCR agree with those by RNA sequencing (Fig. 6), considering a Pearson correlation coefficient > 0.5, except for *norZ*, where no significant changes in the expression of this gene was quantified (fold-change < 2).

**Figure 6.**
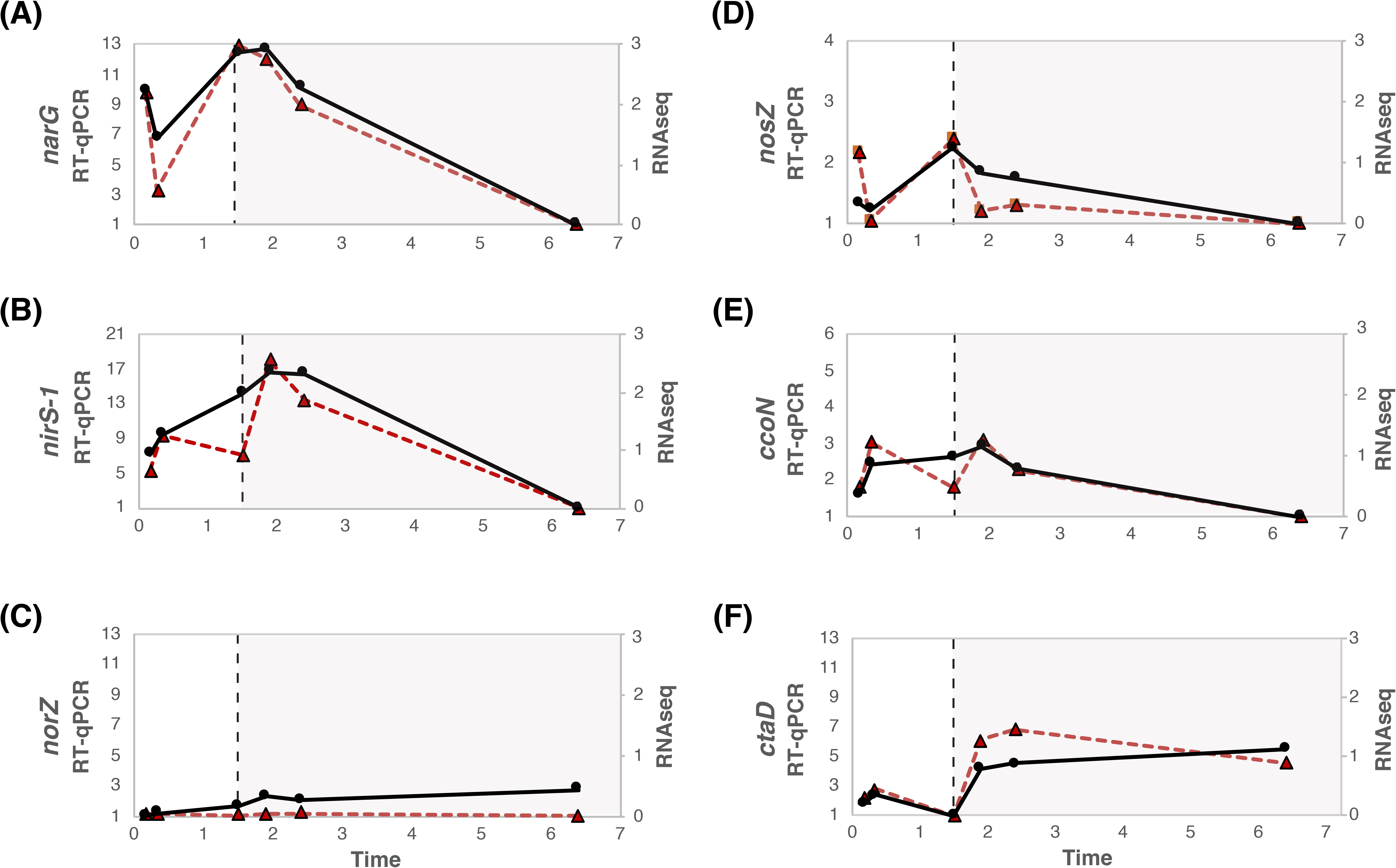
Comparison of the transcriptomic profiles obtained from RT-qPCR (gene copies normalized by *rpoN* copies, solid black lines) and RNA sequencing (ΔLog_2_ RPKM, dotted red lines). Expression of genes involved in denitrification, *narG, nirS, norZ* and *nosZ*, are presented in panels A-D. Genes involved in aerobic respiration, *ccoN* (*cbb_3_*) and *ctaD* (*aa_3_*) are included in panels E-F.

### FNR-type regulator controlling denitrification and aerobic respiration in Accumulibacter

To identify putative regulatory mechanisms of Accumulibacter’s respiratory machinery, an upstream motif analysis was conducted. A sequence motif was identified in the intergenic regions upstream of genes with similar expression patterns: *nar*, *nir*, *nos* and *cbb_3_* operons of Accumulibacter UW-LDO-IC (Fig. 7A). Comparison of this motif with the Prokaryote DNA motif database, using the scanning algorithm Tomtom (73) (p-value = 3.04e-08), classified this sequence as the binding site of FNR, a relatively well-studied member of the CRP/FNR family of transcriptional regulators previously characterized in other proteobacteria (74). FNR is a global regulator of the anaerobic metabolism, reported to be necessary for expression of denitrification and aerobic respiratory pathways (75) and its activity is directly inhibited by oxygen via destruction of a labile iron-sulfur cluster (76).

**Figure 7.**
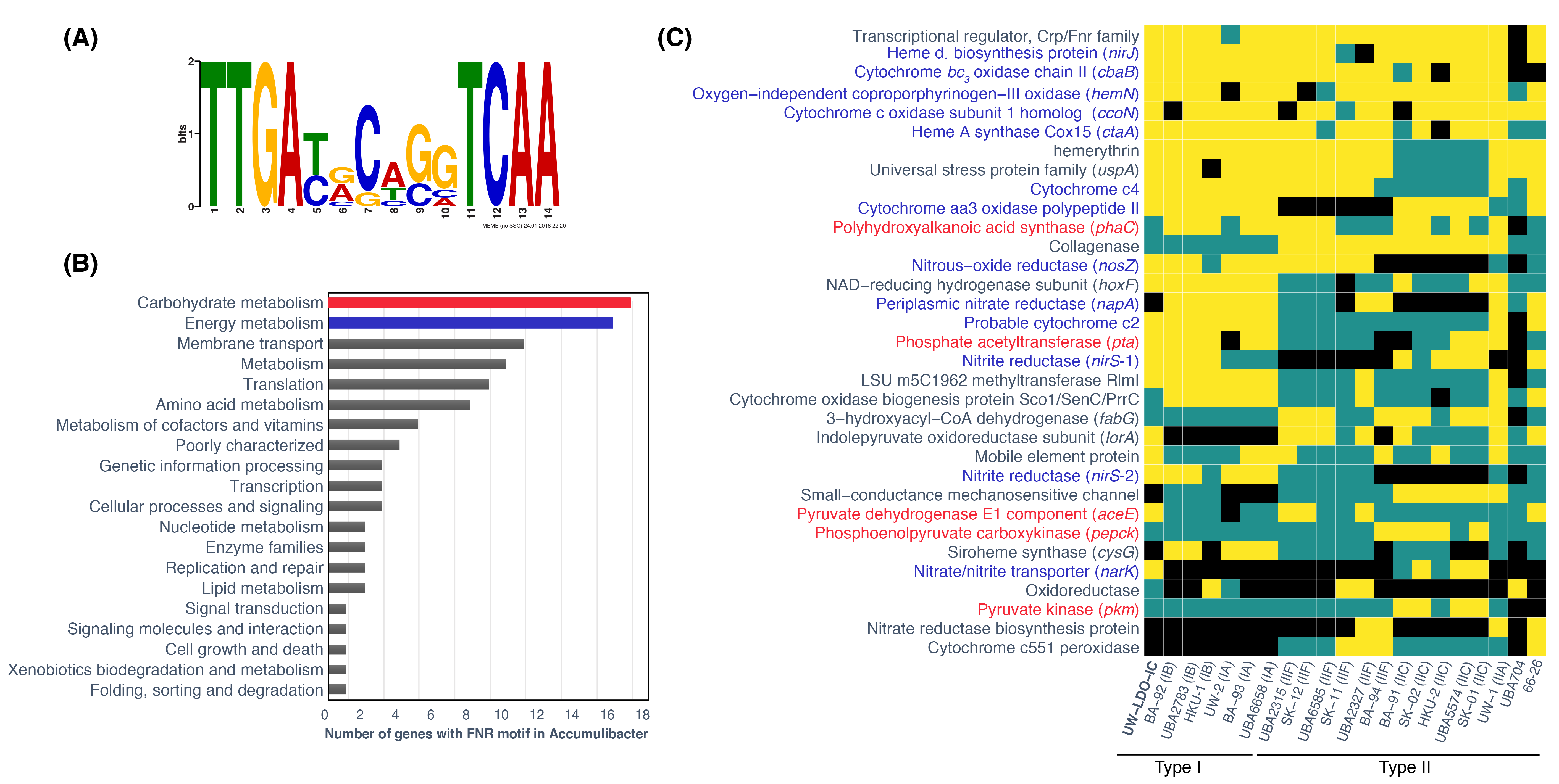
The FNR regulon in Accumulibacter. (A) Motif diagram showing a putative FNR-binding site identified upstream of respiratory genes. (B) Distribution of KEGG functional categories in the fraction of genes with the FNR motif in genomes of Accumulibacter. (C) The predicted conservation of the FNR regulon determined in the Accumulibacter lineage; only genes with motifs present in at least 4 genomes are shown. Black indicates that the corresponding strain does not possess the corresponding gene, blue indicates that the strain possesses the gene, but an FNR-motif was not located within the upstream intergenic region. Yellow indicates that the gene with an FNR-motif is present in the genome.

Subsequent homology searches of this motif sequence were performed in the intergenic region of other genomes of Accumulibacter (p-values < 1e-05). The computational analysis predicted this motif to be located upstream of 165 different genes/operons across all genomes. According to the KEGG category III classification, the majority of these genes appear to be part of metabolic processes involved in carbohydrate and energy metabolism (Fig. 7B). Fig. 7C lists genes with an FNR motif present in at least four Accumulibacter genomes. According to this analysis, FNR auto regulates its own expression, the transcription of genes involved in (a) denitrification, (b) aerobic respiration, including the *aa_3_* and *ba_3_* cytochrome oxidases operons, (c) the biosynthesis of tetrapyrrole heme rings, a prosthetic group of many proteins involved in respiration and the metabolism and transport of oxygen (77), and (c) cytochromes c2 and c4, which are electron donors for *aa_3_* and *cbb_3_* oxidases (78, 79).

Furthermore, FNR seems to participate in the regulation of carbon uptake, since in many cases, its binding site was positioned upstream of a phosphate acetyltransferase (*pta*) (Fig. 7C). Previously, another palindromic sequence was identified upstream of this gene and postulated this motif as the transcriptional factor PhaR binding site (46). Our findings point to another regulatory mechanism for carbon uptake, relying on oxygen availability. In *Escherichia coli*, chromatin immunoprecipitation sequencing (ChIP-seq) tests revealed a putative binding site for FNR upstream of the *pta* operon (80) and RT-qPCR experiments in *fnr* mutants demonstrated that this regulator has a positive effect on the operon transcription (81). Although the FNR binding site was also located upstream of polyhydroxyalkanoic acid (PHA) synthase (*phaC*) in multiple genomes of Accumulibacter, S. M. He and K. D. McMahon (69) did not observe changes in the transcript abundance of *phaC* after exposure to oxygen and, to our knowledge, FNR regulation has not been connected to PHA synthesis in other studied organisms. However, since regulation may vary among Accumulibacter clades, further experiments still need to be carried out to evaluate the effect of oxygen on PHA accumulation in Accumulibacter. Furthermore, in a few Accumulibacter genomes, the FNR motif was also found upstream of genes involved in glycolysis/gluconeogenesis (pyruvate dehydrogenase E1 component (*aceE*), phosphoenolpyruvate carboxykinase (*pepck*) and pyruvate kinase (*pkm*)), which may indicate evolution of this pathway towards an oxygen-independent mechanism. Overall, these results provide evidence for oxygen-driven gene expression regulation, not only as an important factor in the adaptation of Accumulibacter’s metabolism to low-DO and anoxic conditions, but also in that oxygen concentration may directly influence Accumulibacter’s ability to metabolize carbon.

The transcriptional pattern of genes with putative FNR binding sites in UW-LDO-IC were analyzed for O_2_-dependant changes in transcript abundance. The expression profile of 25 operons were grouped by similarity (Pearson correlation), resulting in 6 clusters with different transcriptional patterns regulated in an oxygen-dependent manner (Fig. 8). Three of these clusters were associated with positive control by FNR, since they showed increase in transcript abundance during early anaerobic (Cluster A), late anaerobic (Cluster B) and early micro-aerobic (Cluster C) stages. These three clusters included genes implicated in denitrification (*nar*, *nir* and *nos* operons), micro-aerobic respiration (*cco* operon and *hemN*), electron transport (cytochrome c2 and c4), response to stress (*uspA*), oxygen detection (hemerythrin) and acetate-uptake (*pta*). On the contrary, Clusters C, D and E showed negative regulation by FNR (Fig. 8). The unique transcriptional profile of the transcriptional regulator FNR (Cluster C) might be attributed to the effect of negative autoregulation that FNR has on its own transcription during anaerobic conditions (82). Finally, Clusters D and E were induced during the aerobic stage, with Cluster D comprising genes induced during the entire aerobic stage, and Cluster E genes with higher expression at the end of the cycle (Fig. 8). The machinery for aerobic respiration under high-oxygen conditions was enclosed by Cluster E, including the low-O_2_ affinity cytochromes *aa_3_* and *ba_3_* oxidases and a protein involved in the synthesis of heme A, a prosthetic group required by cytochrome a-type respiratory oxidases. Interestingly, the first subunit of the *ba_3_* oxidase enzyme was not identified as part of this operon, which may explain the differences observed in the expression levels (Fig. 5C). A second copy of the nitrite reductase enzyme (*nirS-2*) also clustered within this group displaying higher transcript abundance during the aerobic stage. Differences in the regulatory mechanisms of these homologs might be a consequence of gene acquisition from different systems.

**Figure 8.**
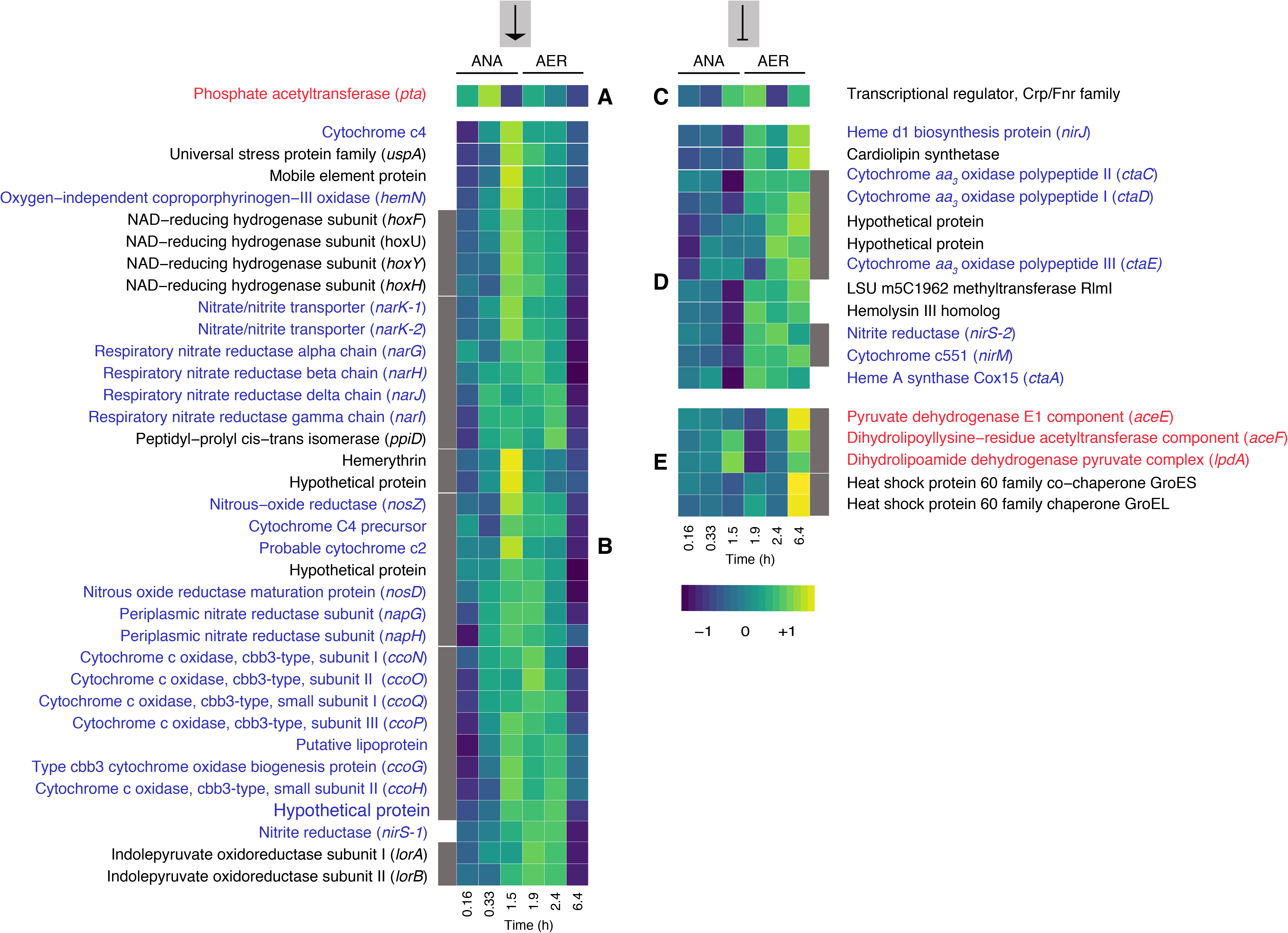
Transcription profile heatmap of members of the FNR regulon in UW-LDO-IC. The colors represent the relative level of mRNA abundance at each time point of the cycle compared to the mean level of expression (yellow=high expression, dark blue=low expression). Genes were clustered according to their expression profiles. Lateral grey bars indicate genes belonging to the same operon. Group A and B contain genes whose expression levels negatively correlate with oxygen tension (positively regulated by FNR). Group C, D and E contains genes upregulated during high oxygen levels (negatively regulated by FNR). ANA: anaerobic, AER: micro-aerobic.

The existence of different expression patterns in genes putatively regulated by FNR, indicates that its transcription could be affected by additional regulatory mechanisms, such as combinatorial binding of other regulators and/or epigenetic signals (83). Other transcriptional factors identified in the Accumulibacter genomes, including NreABC, NsrR, NorR and RegAB, might also serve as signals of none or low-oxygen concentration and regulate part of the anoxic and aerobic respiratory pathways, as it has been described in other microorganisms (84-87). However, at the time, no other binding site associated to one of these regulatory elements has been identified in Accumulibacter.

Overall, this study dissects the metabolic response of Accumulibacter to oxygen-limited conditions. The comparative genomic results provide evidence for the unique respiratory machinery encoded in the newly assembled genome, Accumulibacter UW-LDO-IC, which confers this strain the capability to simultaneously reduce oxygen and nitrogenous compounds. Simultaneous upregulation of both aerobic and anoxic respiratory pathways and co-regulation by the FNR transcriptional factor further support the use of multiple electron acceptors by UW-LDO-IC. Further studies should include experiments analyzing the transcriptional regulation of these pathways at a genome-scale level, using modern approaches such as transcriptional regulatory networks (TRN) and genome wide binding site-locations methods, like ChIP-seq (88) and DNA affinity purification (DAP)- sequencing (89).

## ACKNOWLEDGMENTS

This work was partially supported by funding from the National Science Foundation (CBET-1435661, CBET-1803055 and MCB-1518130), the UW-Madison Office of the Vice Chancellor for Research and Graduate Education, and the Madison Metropolitan Sewerage District. Additional funding from the Chilean National Commission for Scientific and Technological Research (CONICYT) as a fellowship to Pamela Camejo is also acknowledged. The work conducted by the U.S. Department of Energy Joint Genome Institute, a DOE Office of Science User Facility, is supported by the Office of Science of the U.S. Department of Energy under Contract No. DE-AC02-05CH11231. Any opinions expressed in this paper are those of the authors and do not necessarily reflect the views of the agency; therefore, no official endorsement should be inferred. Any mention of trade names or commercial products does not constitute endorsement or recommendation for use.

